# Assessing the sex-related genomic composition difference using a K-mer-based approach: a case of study in *Arapaima gigas* (Pirarucu)

**DOI:** 10.1101/2020.03.29.014647

**Authors:** R.L.D Cavalcante, J.M. Ortega, J.E.S Souza, T. Sakamoto

## Abstract

*Arapaima gigas* is one of the largest freshwater bony fish in the world, in which adults could weigh 200 kilograms and measure 3 meters in length. Due to its large size and its low-fat meat, *Arapaima gigas* has quickly become a species of special interest in fish-farming. One challenge faced during their production is the lack of an efficient sexing methodology, since their sexual maturation occurs late (around the third to the fifth year) and the genetic mechanisms linked to their sex determination system are not known yet. For a more sustainable management, it is of paramount importance to seek an effective and non-invasive method to differentiate sexually juvenile individuals of *Arapaima gigas*. For this, the establishment of genetic markers associated with sexual differentiation would be an advantageous tool. In this study, we proposed a k-mer based approach to identify genome features with sex-determining properties. For this purpose, we used genomic data from four adult representatives of *Arapaima gigas*, two males and two females, and counted the k-mers comprising them. As result, we found k-mers from repetitive regions with high difference and disproportion in the count among individuals of the opposite sex. These differences in the k-mer-based genomic composition indicate the existence of genetic factors involved in the sexing of individuals in *Arapaima gigas*.

## BACKGROUND

*Arapaima gigas*, commonly known as “Pirarucu” or “Paiche”, is the largest bony freshwater fish in the world. It belongs to the bonytongues (Order Osteoglossiformes) Arapaimidae family, and has a natural habitat in the Amazon Basin [1]. Adult specimens may weigh around 200 kilograms and measure about 3 meters [2, 3]. Due to its large size, its flesh containing low-fat and low fishbone, along with its physiological characteristic of emerging to the surface at intervals of 15 minutes to assimilate oxygen, *Arapaima gigas* became a vulnerable species to overfishing in the Amazon region [4] leading to greater surveillance of the marketing of Pirarucu by the Brazilian government in the early 2000s [5, 6].

Some studies show that the use of Pirarucu in intensive fish farming is facilitated, in part, by the physiological characteristics of the animal that guarantee the rusticity of the species [7]. For example, the obligate air-breathing causes this species can tolerate environments with low concentrations of dissolved oxygen in the water [8]. In addition to the facility for captive management, attributes such as the low content of fat, combined with the rapid growth of the species, which weighs an average of 10 kg in its first year of life, add value for an intensification in the commercial exploitation of the *Arapaima gigas* [5, 9, 10].

One of the problems related to its fishing exploitation and fish-farming is that we do not know for certain the genetic mechanisms linked to sex-differentiation in *Arapaima gigas* [11], its sexual maturation occurs around the third to the fifth year of life, and sexual dimorphism is not a strong feature of the species [10]. In recent decades, the creation of *Arapaima gigas* in captivity has been increasingly stimulated, either to develop research to better know the particularities of the species or to exploit its economic potential [12, 13]. For more sustainable management, it is of paramount importance to seek an effective and non-invasive method to differentiate sexually juvenile individuals.

For this, the establishment of a molecular genetic marker related to sexual differentiation would be an advantageous tool. Previous analysis of the *Arapaima gigas* genome found no genes associated with the identification of the sex determination system of these individuals [14, 15]. And chromosomal characterization studies could not distinguishable cytologically a sex chromosome in *Arapaima gigas* [16, 17]. In this study, we proposed to asses the genomic composition of *A. gigas* using a k-mer-based approach to identify regions in excess or missing in one of the sexes.

## MATERIALS AND METHODS

### *A. gigas* sequencing data and data processing

For the following analyses, genomic data were used from four adult *Arapaima gigas* representatives, two males (M1 and M2) and two females (F1 and F2). All samples were collected from Bioproject PRJEB22808 available in National Center for Biotechnology Information (NCBI) database [14]. The quality of the reads was verified with the help of FastQC (v. 0.11.4) [18], and low-quality reads were trimmed with the help of the Sickle paired-end (v. 1.33) [19].

### K-mer Analysis

The K-mer analysis was performed in three steps: (1) k-mer count, (2) k-mer count normalization and (3) k-mer count comparison. The k-mer count was performed with the help of the tool Alignment and Assembly Free (AAF, v. 20171001) [20], a free software written in Python (v. 3.7.0) whose premise is to assemble a phylogenetic distribution based on the k-mers count shared by the samples. The trimmed fastq files of the 4 representatives *Arapaima gigas* were submitted as input data to the script aaf phylokmer.py of AAF using the parameter -k (k-mer size) as 23. The output of this script is a table with n x m elements, in which n is the total number of distinct k-mers (i = 1, 2, …, n), m is the number of samples (j = 1, 2, …, m) and C_ij_ is the count of k-mer i in sample j. For further analysis, we only considered k-mers in which the sum of the count in all samples were greater than 5.

Since the genomic data submitted to this analysis have different coverage values, in order to normalize the data and to decrease the number of false positives, the method Quantile Normalization (QN), a global adjustment method, which consists of a non-parametric methods that makes two or more distributions identical on statistical properties [21].

Finally, to verify the difference in the k-mer abundance among the samples, we compared the samples in pairwise manner (six comparisons in the total) and calculated, for each k-mer, (1) the difference (D_i,j x j’_ = C_ij_ - C_ij’_) and (2) the logarithm of the ratio (L_i,j x j’_ = log2(C_ij_ / C_ij’_)) of their count values. To select those k-mers that showed high D_i,j x j’_ and extreme values of L_i,j x j’_, we calculated the product of both values (P_i,j x j_ = D_i,j x j_ x L_i,j x j_) and extracted the 500 k-mers with highest product value. In the end, 3000 k-mers were extracted from all six comparisons and submitted to CD-HIT tool (v. 4.7) [22] for cluster analysis. Clusters with k-mers that were extracted from a comparison between samples of the same sex was discarded for further analysis. Then, in the remaining clusters, those which contain k-mers extracted from all comparison between samples of opposite sex were considered as candidate genetic markers for sex determination. All graphic plots were generated using Matplotlib (v. 3.0.1) [23] and Seaborn (v. 0.9.0) [24], which are Python 2D plotting libraries.

## RESULTS

The genome sequencing data of 4 samples of *A. gigas*, two female (F1 and F2) and two male (M1 and M2) samples, was obtained and processed. After trimming for low-quality sequences, 8.8 million, 37.7 million, 37.1 million, 13.5 million reads remained for F1, F2, M1 and M2, respectively. The remained reads was submitted to the software AAF [20] to count the k-mers comprising each genome samples. The total k-mers counted was 1.03 billion, 2.15 billion, 2.17 billion and 1.18 billion k-mers for F1, F2, M1 and M2 samples, respectively. Because of the difference on the sequencing depth of each sample (Figure 1A), the Quantile Normalization method (QN) was applied to make the counts comparable between samples (Figure 1B).

**Figure 1.**
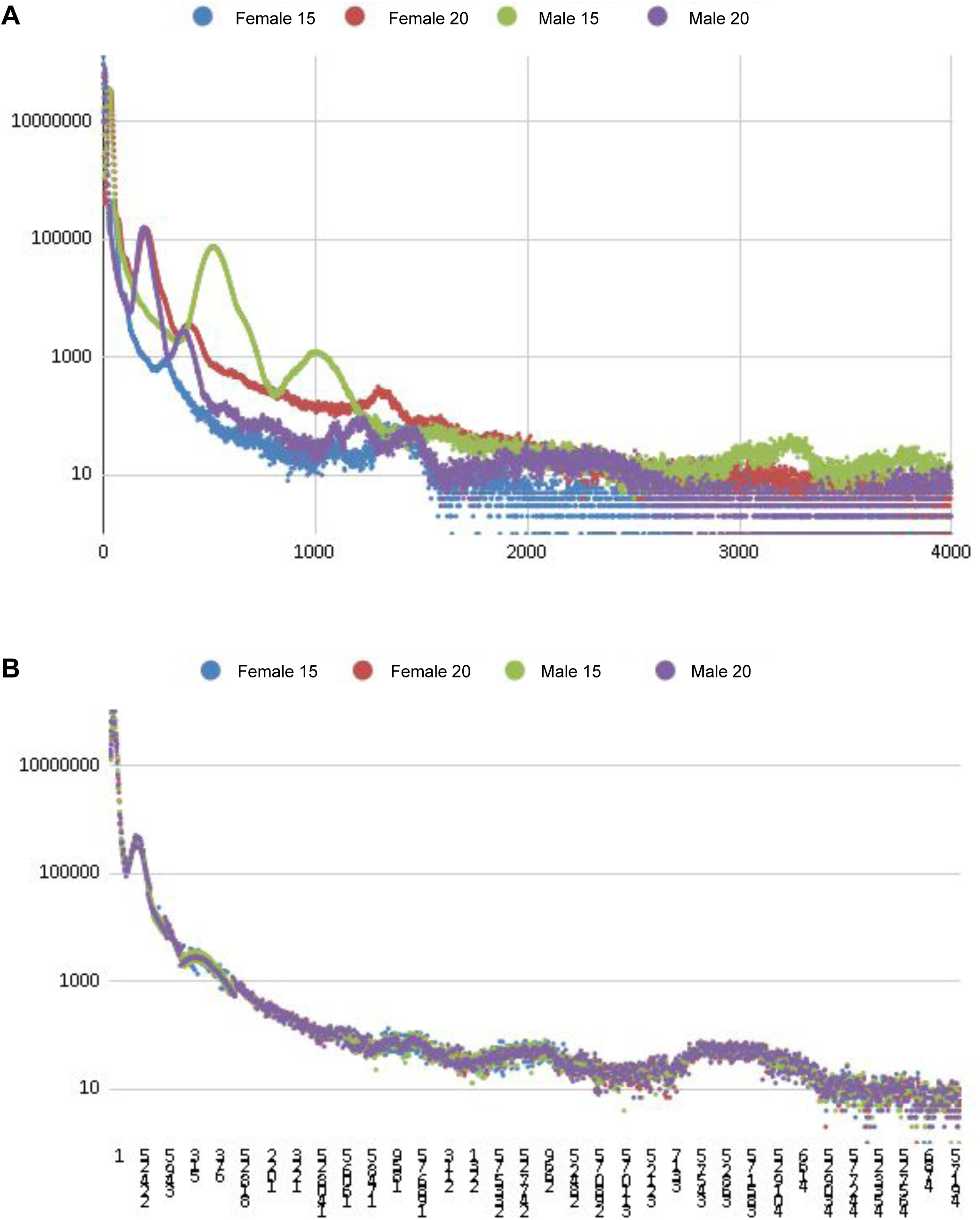
K-mers count size distribution. The distributions were generated using raw (A) and QN normalized (B) k-mer count values. X-axis represents the k-mer count size and Y-axis represents the number of k-mer with that size in log scale.

The comparison of k-mers count between individuals of the opposite sex and the same sex, either by the count difference or logarithm of the ratio, showed some k-mers which were in excess in one of the samples (Figure 2). In each comparison between a pair of samples (6 comparisons in total), we selected 500 k-mers that showed the highest value of the product of the count difference and the log ratio. All selected k-mers (6 × 500 k-mers) had their sequences clustered with CD-HIT [22]. This analysis resulted in the formation of 1129 clusters, of which 104 were comprised of k-mers that was selected only in comparisons between samples of opposite sex. Furthermore, of these 104 clusters, four of them had at least one representative of each comparison made between samples of opposite sex, which were comprised of k-mers of a repetitive regions (Figure 3).

**Figure 2.**
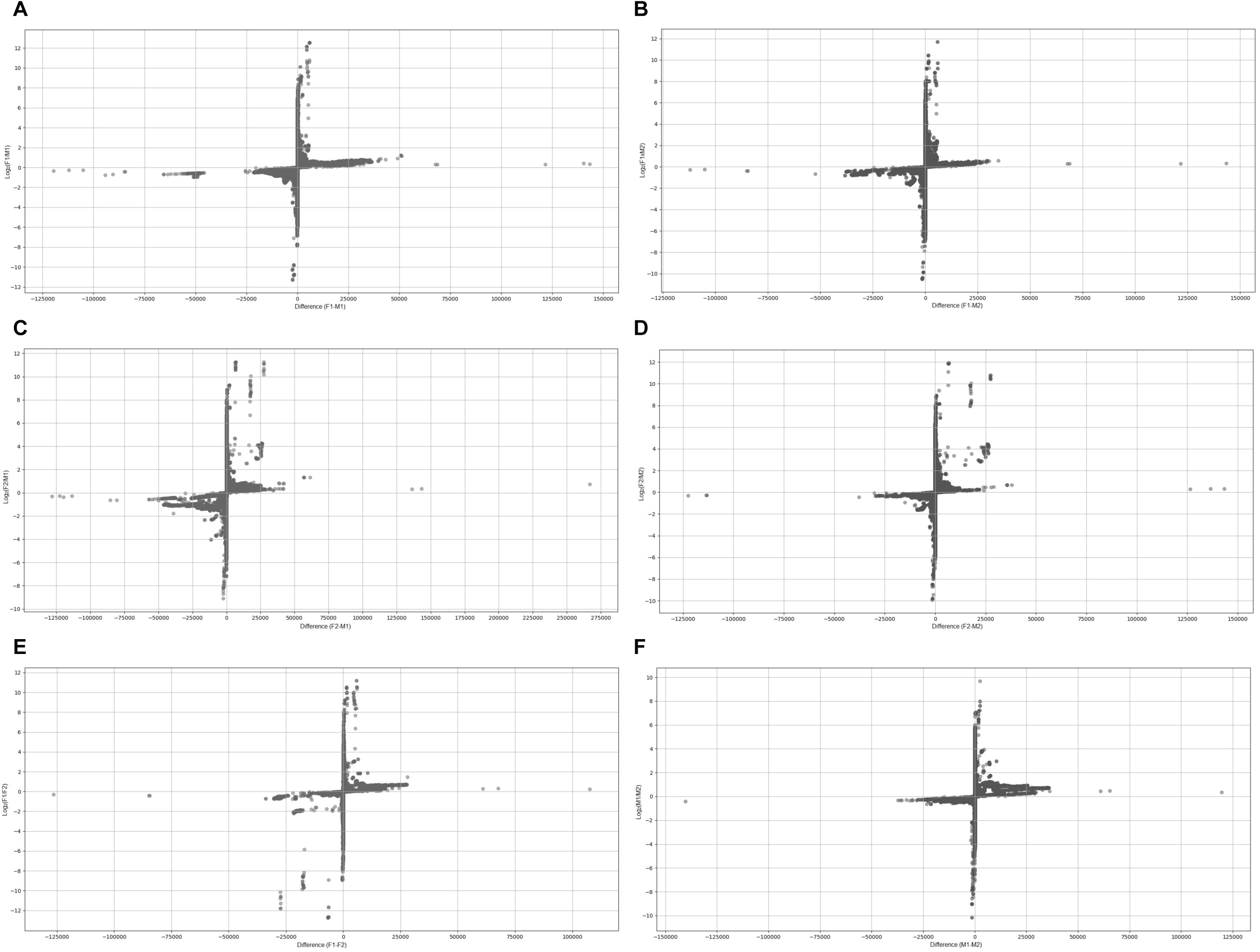
Scatter plot of the k-mer count difference and log ratio between two sample. The k-mer count comparisons between samples F1xM1 (A), F1xM2 (B), F2xM1 (C), F2xM2 (D), F1xF2 (E) and M1xM2 (F) are shown. For all plots, each dot represents a k-mer, X-axis represents the k-mer count difference and Y-axis represents the log ratio of the k-mer count dots far from the origin represent k-mers in excess in one of the sample.

**Figure 3.**
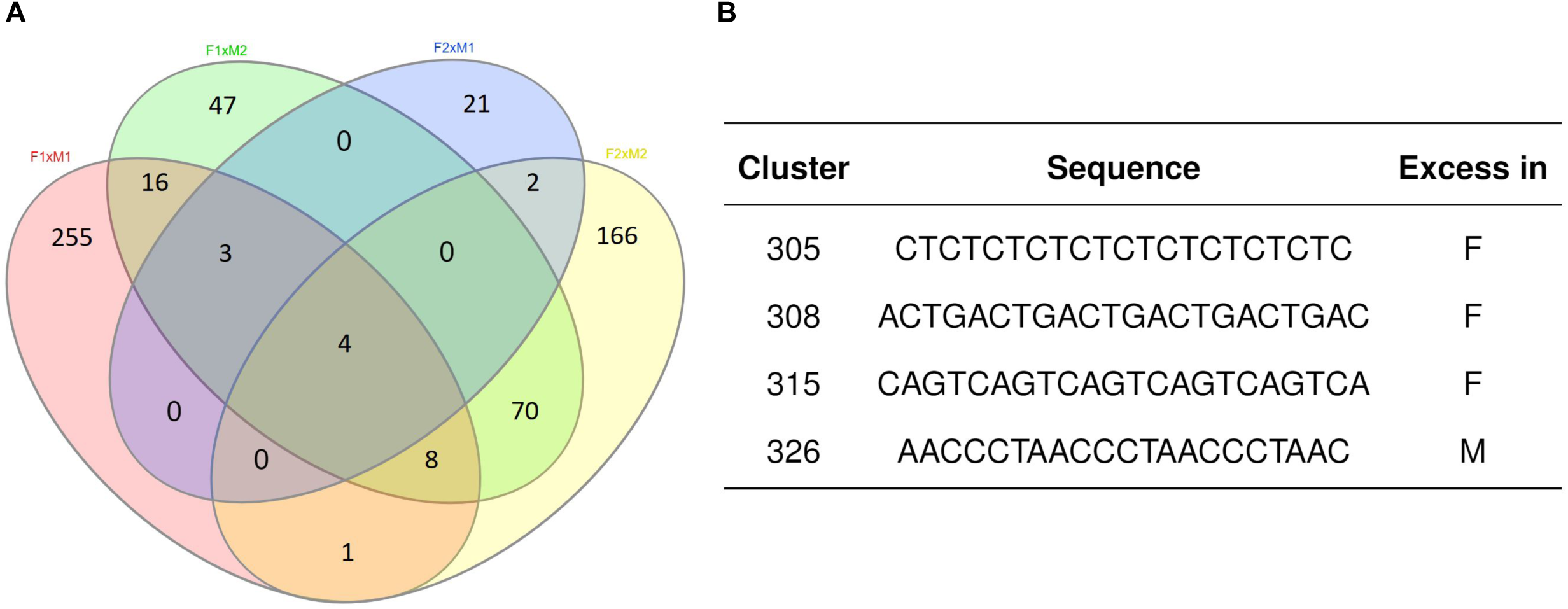
The 104 clusters comprised of k-mers extracted only from the comparison between individuals of opposite sex. A) Venn Diagram showing the k-mers shared among those extracted from the comparison F1xM1, F1xM2, F2xM1 and F2xM2. Four clusters showed are in the intersection of all comparison. B) Table detailing the four clusters in the intersection. F: female; M: male.

## DISCUSSION

Understand the mechanisms of sexual determination in fish are essential for a sustainable management of ichthyofauna, either for commercial or conservation purposes [25]. Introspection of these mechanisms in *A. gigas* has an appeal because this species plays an important role in the economy of the north region of Brazil [12, 13]. But elucidating these mechanisms in fish is challenging, since most of them, including *A. gigas*, do not bear a typical sex chromosome in their genome [16, 17] and the difference of the genome sequences between samples of opposite sex seems to be minimal. The first genome sequencing of *A. gigas* [14] did not find significant differences between the genomic content of male and female samples. The second one [15] suggested the XY system for *A. gigas*, but their results conflict with the genome data of the first one.

The approach of finding specific regions in one sex using genomic data has demonstrated to be not so effective in finding those regions associated with the sex-determination. In this context, we explored other genomic features in *A. gigas* to find some clues about the genetic factors involved in the sex determining system. For this, we analysed the genomic composition of Pirarucu using k-mer based approach.

In this study, we have noticed the existence of k-mers over or underrepresented in one of the sexes, indicating potential differences in the genetic composition between males and females of *A. gigas*. The difference is not so expressive, which corroborates with the reports that estimate 0.01% [14] to 0.1% [15] of the genome of this species as linked to the sexual determination. The four sequences reported in this work (Figure 3) are all part of repetitive sequences. Despite of their low complexity, repetitive regions have been reported to have important role on sex determination [26]. In medaka, which has the XY system, there is a large stretch of repetitive regions on the Y-specific regions [27]. The chromosome Y of Pacific salmon bears a specific repetitive regions (OtY1) that is used as genetic marker to differentiate sex [28].

In this context, the repetitive sequences found in this study could be a component that could be used to determine sex in individuals of *Arapaima gigas*. We recognize, however, the necessity to perform analyses with a greater number of samples to obtain a statistical support for our results. Despite of that, the kmer-based method applied on this work has demonstrated to be an interesting strategy to help us discover the sex-determination system in Pirarucu specimens and can be extended to other species.

## CONCLUSIONS

With this study, we were able to suggest some repetitive genome sequences that may be differentiated in quantity in male and female of *A. gigas*. We consider these regions as strong candidates for a molecular marker that could be used for sexing individuals of *A. gigas*. However, because of the small sample size, new analyses with a larger number of individuals are necessary to allow statistical support for our conclusion, as well as to suggest bench trials for the validation of the in-silico analyses. Furthermore, kmer-based methods demonstrated to be an interesting strategy to assist us in unraveling the sex-determination system in Pirarucu.

## ACKNOWLEDGMENTS

This work was supported by CAPES (Coordenação de Aperfeiçoamento de Pessoal do Ensino Superior). To the Instituto Metrópole Digital - IMD, and specifically to Bioinformatics Multidisciplinary Environment - BioME for the computational structure for the accomplishment of this work. To the Federal University of Rio Grande do Norte - UFRN for structural support and resources have been ceded. And to Dr. Sidney Emanuel Batista dos Santos from Federal University of Pará - UFPA for contributing in discussions in this project.

## AUTHOR CONTRIBUTIONS

RLDC designed the pipeline and evaluated the data. JMO assisted with computational structure. JESS assisted with project development and discussions. TS designed the project, evaluated the data and assisted with project development and discussions.

